# Differential disease tolerance mediates sex-biased illness severity in sepsis

**DOI:** 10.1101/2025.07.28.667282

**Authors:** Breenna Dobson, Kathryn Strayer, Ayesha Wijesinghe, Jared Schlechte, Diana Changirwa, Nicole A Cho, Ian-ling Yu, Braedon McDonald

**Author notes:** Corresponding Author: Braedon McDonald, MD, PhD, FRCPC Associate Professor Department of Critical Care Medicine Snyder Institute for Chronic Diseases Cumming School of Medicine, University of Calgary 3330 Hospital Dr NW, Calgary, AB, Canada T2N 4N1 1-403-220-6885.

## Abstract

Sepsis in humans, as well as mouse models of infection, demonstrates sex-biased outcomes in which males tend to have a higher incidence, higher severity, and higher mortality compared to females. Despite this important sex-bias in sepsis outcomes, little is known about its mechanistic drivers nor therapeutic implications, as much of the foundational data on sepsis pathogenesis is derived from animal studies that included only male subjects, potentially contributing to the notable paucity of successful mouse-to-human translation of sepsis therapeutics. In this study, we demonstrate that male-biased illness severity and organ dysfunction in mouse models of bacterial sepsis is mediated by impaired disease tolerance in males, involving impaired tolerogenic shifts in mitochondrial oxidative metabolism compared to females. Microbiological analyses and systems immunology characterization of sepsis between males and females revealed that sex-biased disease tolerance was independent of infection resistance mechanisms, as well as canonical immune/inflammatory dysregulation. Therapeutic potentiation of mitochondrial tolerance with doxycycline neutralized sexual dimorphism of illness severity and organ dysfunction through a male-predominant treatment effect. These data reveal that biological sex is a fundamental determinant of illness severity and treatment-responsiveness in sepsis through modulation of disease tolerance, which may be harnessed therapeutically to address sex-biased outcomes in sepsis.

**One sentence summary:** Sex-biased illness severity in bacterial sepsis is caused by impaired disease tolerance in males that can be rescued by therapeutic potentiation of mitochondrial tolerance

## INTRODUCTION

Studies of human sepsis epidemiology as well as pre-clinical animal models of severe infections have identified sex-biased outcomes in which males experience a higher incidence, higher illness severity, and higher mortality compared to females(*1–9*). The underlying mechanisms that mediate this sex bias in sepsis outcomes are unknown, in part because many foundational discoveries in the field of sepsis research have been derived from animal models that included only male subjects(*10–12*). As such, inappropriate generalizations about sepsis pathogenesis based on unrecognized sex-dependent mechanisms may contribute to the near-universal failure of (male) mouse-to-human translation of therapeutic discoveries in human clinical trials(*13*).

Therefore, determining mechanisms that mediate sex-biased outcomes in sepsis may reveal impactful opportunities to re-examine previous “failed” sepsis therapies through a sex-inclusive lens, and develop novel precision-based approaches to treatment that account for differential disease mechanisms and/or treatment responsiveness between males and females.

Sepsis is life-threatening organ dysfunction caused by a dysregulated host response to infection, which in its severe form (septic shock) is associated with 25-30% mortality(*14*). The mechanisms of dysregulated host response to infection that cause sepsis broadly fall into 2 categories – dysregulated infection *resistance* (failure of host mechanisms that eradicate infecting pathogens), and dysregulated infection *tolerance* (failure of mechanisms that protect host cells and tissues from damage while combatting infection)(*15*). The pathogenesis of sepsis reflects the classical “double-edged sword” between disease resistance and tolerance. An effective immune response is needed to protect the host from overwhelming infection, but this occurs at the expense of damage to host cells and tissues leading to organ dysfunction that is the defining feature of sepsis(*16*)(*17*). More recently, multiple studies have uncovered a crucial role for mechanisms of disease tolerance involving mitochondrial stress responses to infection, termed mitochondrial tolerance, that can be harnessed therapeutically to improve outcomes in animal models of sepsis(*18, 19*). However, in keeping with the historical trend in preclinical sepsis research, studies of mitochondrial tolerance and its therapeutic potential in bacterial infections were performed in male mice only (*18, 19*). Therefore, the impact of biological sex on mechanisms of infection resistance and tolerance in bacterial sepsis, and their implications towards sex-dependent treatment responses, remains a critical outstanding question in the field.

In this study, we utilized well-established models of murine bacterial infections to identify mechanisms underlying the male bias in sepsis severity and outcomes, and its potential therapeutic implications. In response to both polymicrobial and mono-microbial infections, male mice demonstrated higher illness severity, physiological derangement, and end-organ dysfunction compared to age-matched female littermates. Sex-biased outcomes were driven by gonadal sex, but not sex-chromosome-linked gene differences between males and females.

Extensive characterization of pathogen dissemination, systemic inflammation, cellular immunity, transcriptomic responses, and cellular metabolism revealed that male-biased sepsis severity is driven by impaired infection tolerance due to sex-biased mitochondrial tolerance, and was independent of infection resistance or canonical immune/inflammatory dysregulation during severe infection. As such, therapeutic correction of impaired mitochondrial tolerance demonstrated male-biased efficacy, neutralizing differences in sepsis severity between males and females. Collectively, this study identifies differential infection tolerance mechanisms as mediators of sex-biased sepsis severity and outcomes, and establishes biological sex as a crucial variable in both the study of sepsis pathogenesis as well as treatment responsiveness, paving the way for pre-clinical and clinical research into sex-based precision therapies for sepsis.

## RESULTS

### Male biased illness severity and organ dysfunction during acute bacterial sepsis

Illness severity in response to acute bacterial infection was quantified in age-matched male and female mice (littermates) using physiologic, biochemical, and behavioural measures of sepsis severity. Consistent with current Canadian Council for Animal Care (CACC) guidelines, which stipulate that mortality is not an acceptable endpoint in animal research, sepsis severity was measured using these published and validated non-mortality metrics in sub-lethal models of sepsis. First, we quantitatively assessed sickness behaviour during acute sepsis, 6 hours after polymicrobial infection with fecal-induced peritonitis (FIP). Infected male mice were consistently more lethargic, immobile, huddled, piloerected, and less responsive to stimuli compared to age-matched females. Quantification of sickness behaviour using the Murine Sepsis Severity Index (MSSI)(*20*) confirmed significantly higher MSSI in males (MSSI median 11, range 8-20) compared to females (MSSI median 5, range 1-7) (Fig 1A). Next, we evaluated core temperature as a key physiologic indicator of infection severity, wherein the severity of hypothermia is closely associated with the risk of death from infection in mice. Again, male mice experienced a significant decrement in core body temperature at 3 and 6 hours post infection when compared to pre-infection temperature, whereas female mice did not demonstrate a significant change in core temperature during acute infection (Fig 1B). Furthermore, comparing mean core temperatures between males and females confirmed that males experience significant worse hypothermia compared to females in response to acute bacterial sepsis (Fig 1B).

**Figure 1.**
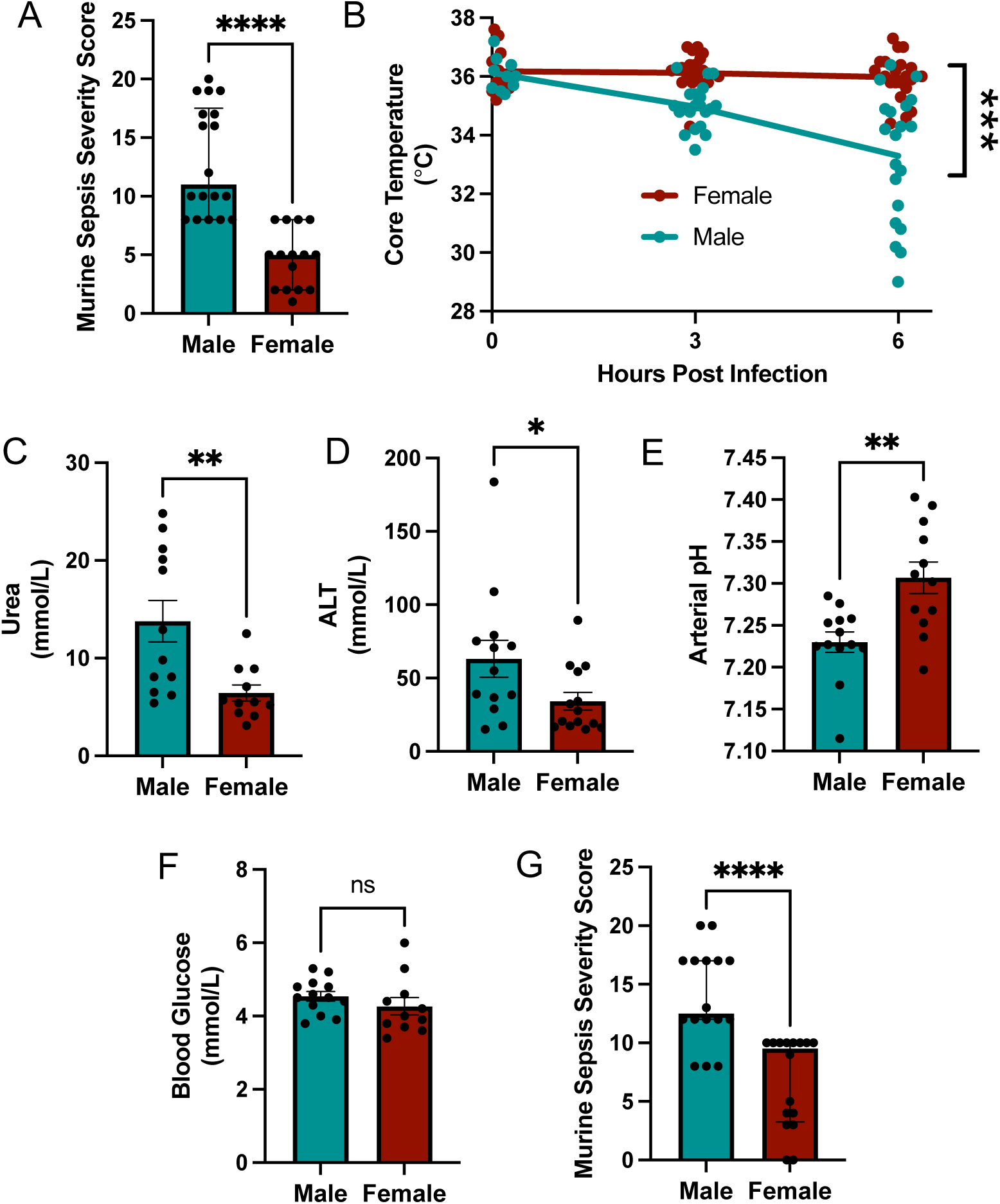
-Male-biased illness severity in murine sepsis. (A) Male (N=18) and female (N=15) age-matched 8-12 week old littermates were infected intraperitoneally with donor cecal contents (1 mg/g) to induce polymicrobial sepsis. Murine sepsis severity index was quantified at 6 hours post infection. (B) Rectal core temperatures were measured at 3 and again at 6 hours post polymicrobial peritonitis infection in males (N=20) and female (N=21) mice. Biochemical measurements of end organ dysfunction including (C) plasma urea, (D) plasma ALT, (E) arterial blood pH, and (F) plasma glucose were quantified in blood collected from male (N=13) and female (N=14) mice at 6 hours post polymicrobial peritonitis infection, with N value varying slightly between assays due to limited sample volume. (E) Murine sepsis severity index was quantified at 6 hours post mono-bacterial infection with *E. coli* Xen14 (2x10^7^ CFU, i.p.). Dots are individual mice, bars show medians +/-interquartile range. Data were analyzed using Mann-Whitney U test, ****p<0.0001, **p<0.01, *p<0.05, ns=p>0.05.

A defining characteristic of sepsis is the development of end organ dysfunction in response to infection(*14*). Biochemical measures of sepsis-related organ dysfunction were quantified in infected male and female mice, including plasma urea (marker of renal dysfunction), plasma alanine aminotransferase (ALT, marker of hepatocellular liver damage), arterial blood pH (measure of metabolic dysfunction), and blood glucose levels. Compared to females, male mice experienced significantly worse renal and liver injury demonstrated by higher plasma urea and ALT levels, as well as more severe acidemia (lower arterial blood pH) (Fig 1C-E). In contrast, no differences were observed in blood glucose levels between sexes (Fig 1F). To determine whether male biased sepsis severity was unique to FIP/polymicrobial sepsis, we repeated these experiments in a mono-bacterial infection (*E. coli* peritonitis), and similarly observed that males experienced significantly higher illness severity compared to females (Fig 1G). Overall, these data support the conclusion that illness severity during acute sepsis is sex-biased, with males experiencing more severe manifestations of illness severity and organ dysfunction compared to females.

### Sex-biased sepsis severity is independent of infection resistance

Higher sepsis severity in males compared to females could be the consequence of differences between sexes in the ability to defend against pathogen growth and dissemination (i.e. infection resistance), or sexual dimorphism of host response to infection (i.e. infection tolerance), or both. To determine whether male-biased sepsis severity could be explained by impaired infection resistance in males, we quantified pathogen burden in mice with fecal peritonitis at 6 hours post infection at the site of infection (peritoneal fluid) as well as sites of dissemination (blood and multiple organs). We hypothesized that sex-biased infection resistance would manifest as increased pathogen burden in males compared to females. Contrary to this hypothesis, we observed no differences in pathogen quantities in peritoneal lavage fluid, blood, spleen, liver, lung, and kidney between males and females (Fig 2A-F). Importantly, high amounts (CFU) of bacteria were recovered in multiple organs from all mice, demonstrating that both sexes had invasive and disseminated infections. However, no differences in the quantity of infecting bacteria were identified, whether cultured under aerobic or anaerobic growth conditions on rich culture media (Supplementary Fig 1). Appreciating the possibility that culture-based quantification of pathogen burden in polymicrobial sepsis may bias for detection of only culturable organism, we repeated this experiment using mono-bacterial infection with *E. coli* and again observed high pathogen burden in all body compartments that was equivalent between males and females (Supplementary Fig 2). Together, these data support the conclusions that that sex-biased sepsis severity cannot be attributed differential infection resistance between males and females, as no differences in pathogen growth and dissemination were observed between sexes in acute bacterial sepsis.

**Figure 2.**
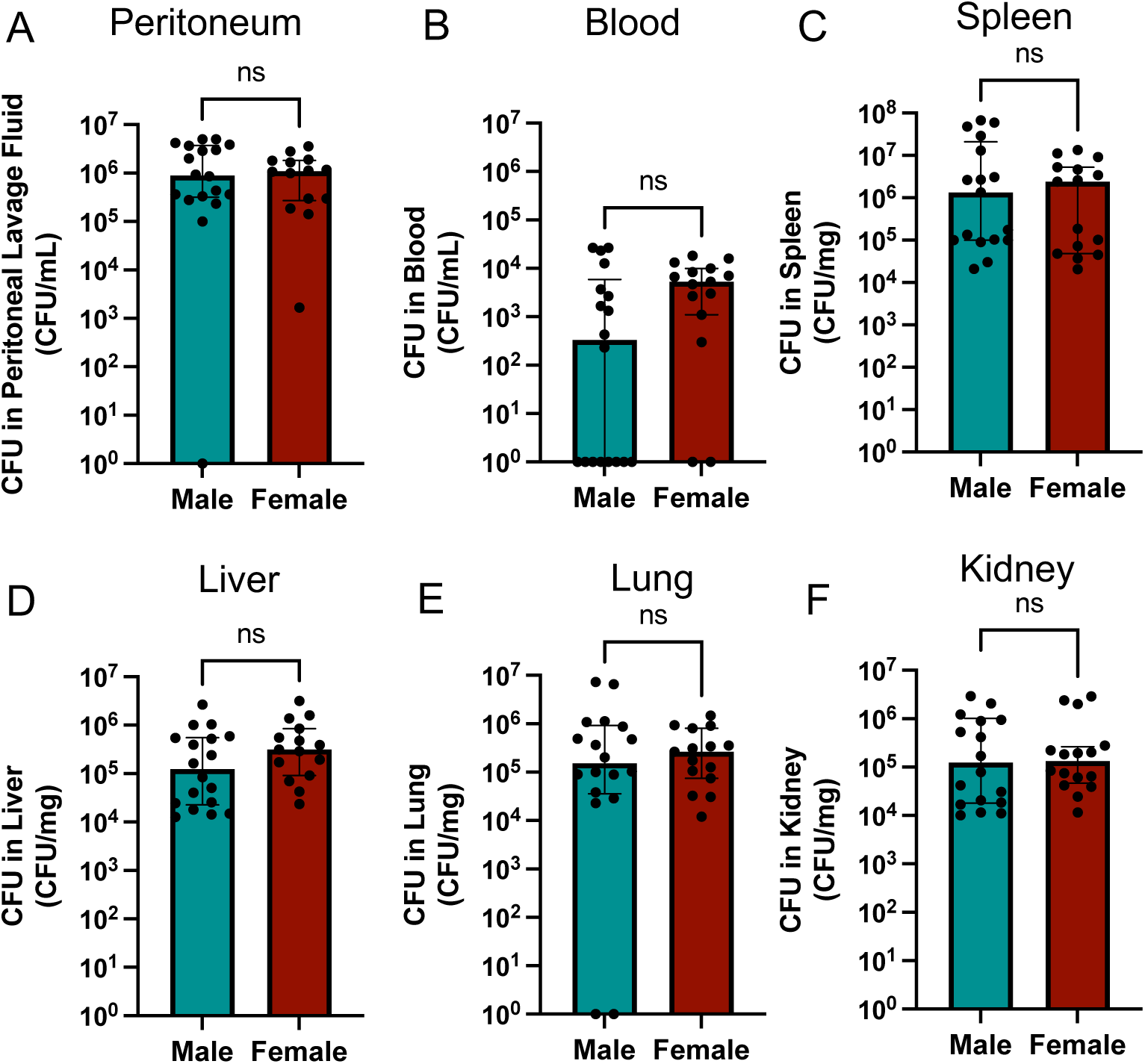
-Equivalent resistance to disseminated infection between males and females with polymicrobial sepsis. Male (N=18) and female (N=15) age-matched 8-12 week old littermates were infected intraperitoneally with donor cecal contents (1 mg/g) to induce polymicrobial sepsis. At 6 hours post-infection, bacterial CFU were quantified in (A) peritoneal fluid, (B) blood, (C) spleen, (D) liver, (E) Lung, and (F) kidney tissue samples. Dots are individual mice, bars show medians +/-interquartile range. Data were analyzed using Mann-Whitney U test, ns=p>0.05.

### Sex-biased sepsis severity is dependent on gonadal phenotype rather than sex-chromosome gene linkage

Sex-biased phenotypes may be the consequence of either sex chromosome-linked genes (e.g. X- or Y-linked genes involved in host response to infection) and/or biological sex phenotypes that may influence host response to infection (gonadal and hormonal composition differences, including testes and androgens in males, vereus ovaries and estrogen/progesterone in females) (*9*). To determine whether sex-biased sepsis severity is dependent on sex chromosome-linked genes versus sex phenotype (gonad/hormone phenotype), we employed a transgenic model that enables uncoupling of sex chromosomes and gonad/hormone phenotype, called the 4 core genotype model(*21*). The 4 core genotypes (XX females with ovaries, XY males with testes, XX males with tested, XY females with ovaries) are generated by breeding wild-type XX females with fertile transgenic XY males in which the sex-determining *sry* gene is deleted from Y chromosome, and an *sry*-transgene inserted into an autosomal chromosome. As a result, sex phenotype (gonads/hormonal composition) in progeny is driven by autosomal *sry* transgene, and thus independent of X and Y chromosomes (Sry^+^ = testes and prototypical male phenotype development, Sry ^-^ = ovaries and prototypical female development), and is independent of sex chromosome composition (Fig 3A).

**Figure 3.**
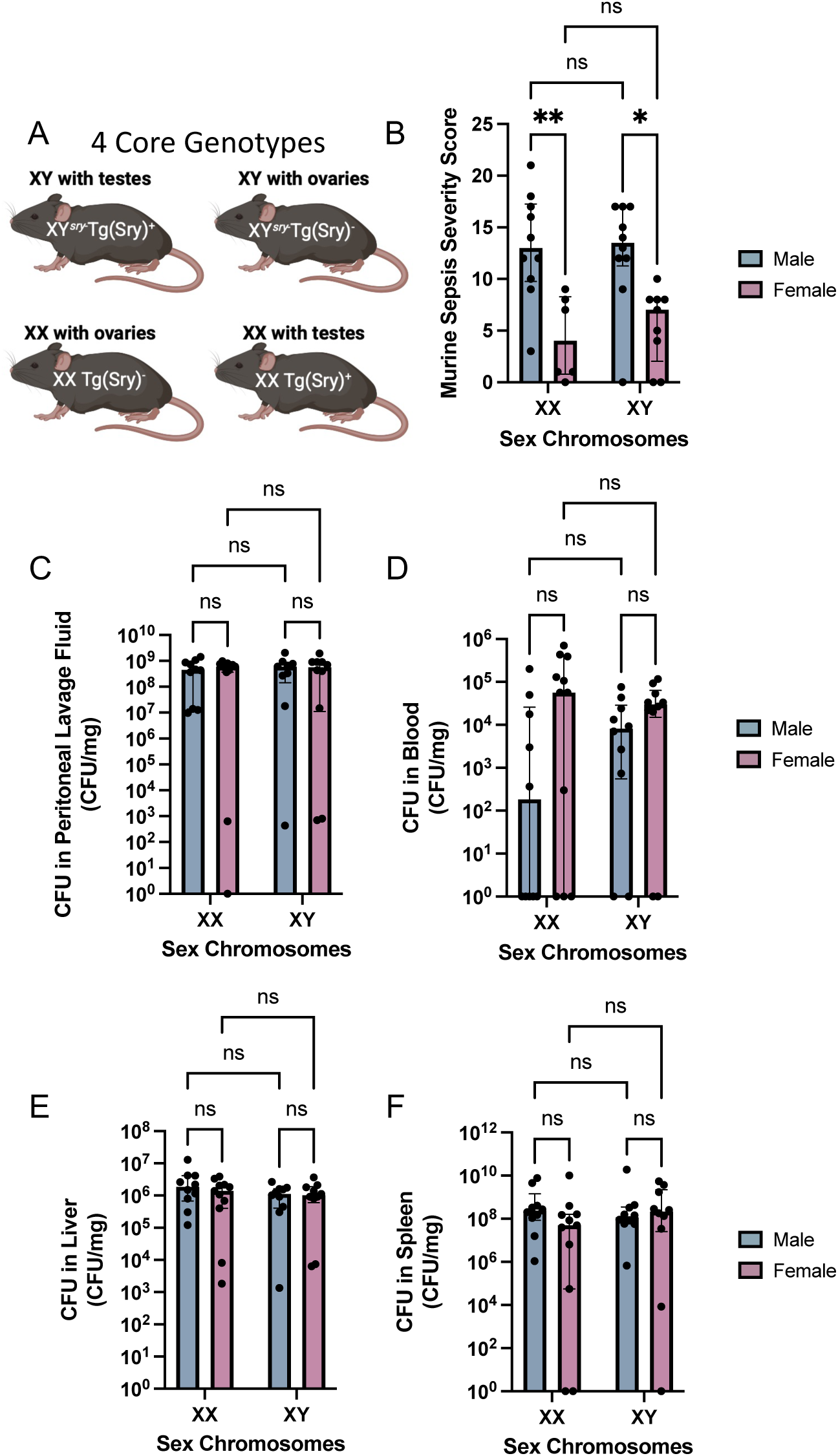
-Sex-biased illness severity in sepsis is mediated by gonadal sex phenotype, not sex chromosomes. (A) Schematic overview of the 4 core genotypes in this transgenic mouse model. (B) Murine sepsis severity index was quantified at 6 hours post infection in XX females (N=6), XX males (N=10), XY females (N=9), and XY males (N=10). Bacterial CFU were quantified in (C) peritoneal fluid, (D) blood, (E) liver, and (F) spleen samples. Dots are individual mice, bars show medians +/-interquartile range. Data were analyzed using Mann-Whitney U test, **p<0.01, *p<0.05, ns=p>0.05.

Sepsis was induced in each of the 4 core genotypes of mice with polymicrobial fecal peritonitis (FIP), and illness severity was quantified using the murine sepsis severity index (MSSI) as above. Interestingly, there was precise demarcation of sepsis severity based on biological sex phenotype, completely independent of sex chromosome composition, in which gonadal males demonstrated significantly higher illness severity during acute sepsis compared to gonadal females (Fig 3B). Again, the observed differences in illness severity were independent of infection resistance, as no differences were observed in pathogen burden across all organs tested between male and female mice (Fig 3C-F). These data suggest that sex-biased illness severity in sepsis is not the result of specific X- or Y- chromosome linked genes, but rather is dependent on gonadal sex phenotype.

### Systemic immune response to sepsis demonstrates limited sexual dimorphism and does not drive sex-biased illness severity

Immune dysregulation and pathological systemic inflammation are considered core components of sepsis pathogenesis. In fact, anti-inflammatory/immunomodulatory therapy with corticosteroids is currently the only guideline-endorsed medical therapy for sepsis aside from early antibiotic administration, intravenous fluids, and physiologic support (*22*). Therefore, we sought to determine whether sex-biased sepsis severity was the result of sexual dimorphism in the systemic immune response to severe infection.

We undertook a comprehensive analysis of the systemic inflammatory response (cytokine storm), as well as cellular immune response in male and female mice in response to acute sepsis. Plasma levels of 16 inflammatory mediators were measured using Luminex immunoassay in the plasma of mice with fecal peritonitis at 6 hours post infection (Fig 4A). 15 of the 16 biomarkers of systemic inflammation were not different between males compared to females, including prototypical pro-inflammatory mediators (IL-6, TNFa, IL-1b, IL-17A, IL-12p70), anti-inflammatory mediators (IL-10, IL-4), and immunomodulatory mediators (IL-2, GM-CSF, IFNg) (Fig 4A). Plasma levels of macrophage inflammatory protein 2 (MIP-2, CXCL2) were slightly higher in males, albeit levels were very high in both males and females. Overall, this evaluation of the systemic “cytokine storm” of severe infection failed to reveal major sexual dimorphism that could explain sex-biased sepsis severity.

**Figure 4.**
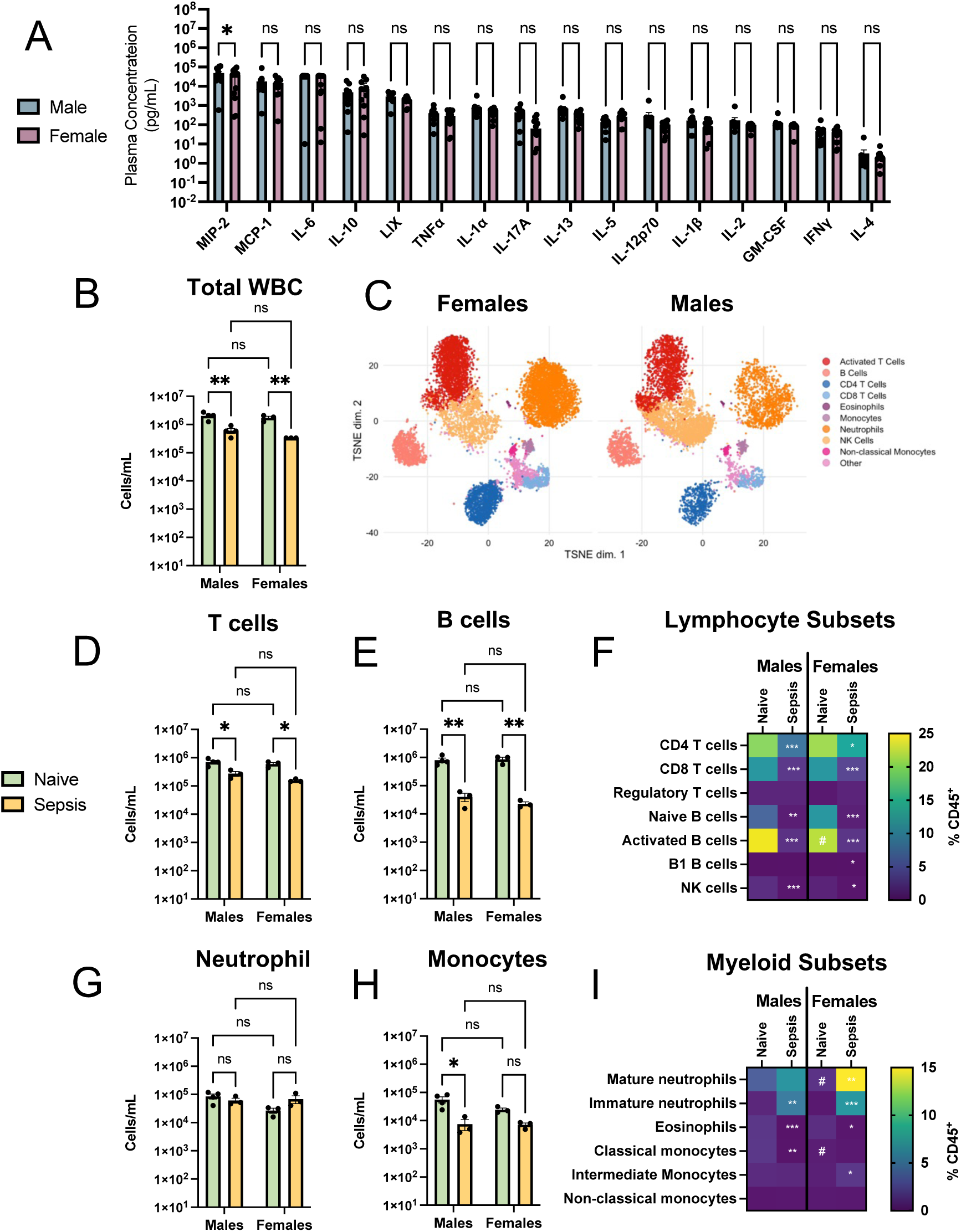
– Minimal differences in systemic inflammation and cellular immune responses during sepsis between males and females. (A) Plasma levels of 16 inflammatory mediators were quantified in the plasma of male (N=12) and female (N=12) mice after 6 hours of polymicrobial peritonitis infection. Mass cytometry analysis of circulating immune cells in the blood was used to quantify various immune cell subsets in both naïve males (N=4) and females (N=3) and septic male (N=3) and females (N=3). (A) Total circulating white blood cells, (B) FLOWSOM clustering of circulating CD45+ leukocytes in males and females, (C) CD45+CD3+ T cells, (D) CD45+CD19+ B cells, (F) multiple lymphocyte subsets (displayed as heatmap of % total CD45+ cells), (G) CD45+CD3-CD19-Ly6G+ neutrophils, (H) CD45+CD3-CD19-Ly6G-Ly6C+ monocytes, and (I) various myeloid cell subsets (displayed as heatmap of % total CD45+ cells). See supplementary figure 3 for gating strategies for each cell population. Dots are individual mice, bars show medians +/-interquartile range. Data were analyzed using 2-way ANOVA, **p<0.01, *p<0.05, ns=p>0.05.

Next, we performed high-dimensional single cell analysis of the systemic immune response in the bloodstream to acute fecal peritonitis. Blood from septic males and females was collected at 6 hours post infection, and analyzed by mass cytometry using a panel of 42 markers to identify and quantify all major immune cell populations in the circulation (Supplementary Table 1, and Supplementary Figure 3). In both males and females, we observed the expected decrement in total circulating white blood cell counts in septic mice as compared to naive (uninfected) controls (Fig 4B). Unsupervised clustering of immune cell populations in the circulation of septic mice using FlowSOM revealed no differences in clustering between males and females (Fig 4C). Within the adaptive arm of the immune response, quantitate analysis of the number of circulating T cells and B cells demonstrated that both males and females experienced equivalent degrees of T and B lymphopenia in response to infection compared to naïve controls (Fig 4D-E). More granular analyses of lymphocyte subsets likewise demonstrated equivalent sepsis-induced reductions in CD4 and CD8 T cell subsets, B cell subsets, an innate lymphoid cells (NK cells) (Fig 4F). Only the rare population of B1 B cells demonstrated slight sex-based differences in response to sepsis, with sepsis-induced changes observed only in females but not males (Fig 4F). Within the innate arm of the immune response, no significant differences were observed in the quantities of circulating neutrophils between males and females, neither naïve nor in response to infection (Fig 4G-H). Within the monocyte compartment, sepsis-induced reductions in circulating monocyte concentrations were observed in males, with a non-significant trend observed in females. Detailed analysis of the relative abundance of neutrophil and monocyte subsets in the circulation demonstrated that females contained relatively higher levels of circulating mature neutrophils and classical monocytes under naïve conditions, whereas other innate cell populations responded similarly to infection in both sexes (Fig 4I). Collectively, minimal differences were observed in the systemic inflammatory and cellular immune response to sepsis between males and females, refuting the hypothesis that higher illness severity in males is driven by sex-biased hyperinflammation and immune dysregulation.

As noted above, immunomodulatory therapy with corticosteroids represents an important tool in the therapeutic armamentarium for human sepsis treatment. In mice, pre-treatment with corticosteroids (hydrocortisone 4 mg/kg, ip) reduced sepsis severity in response to fecal peritonitis compared to vehicle control as demonstrated by significant reduction of median Murine Sepsis Severity Index (MSSI) in treated mice (Fig 5A). To determine whether the observed male bias in sepsis severity translated into differential treatment response to steroid therapy, we compared illness severity, core temperature, and organ dysfunction measures between males and females. In both vehicle control and steroid-treated mice, males consistently demonstrated higher illness severity (MSSI), more severe hypothermia, more severe acidemia, and greater organ dysfunction (Fig 5B-F). However, the impact of steroid treatment on illness severity, thermoregulation, and organ dysfunction was not different between males and females (Fig 5B-F). These results suggest that the therapeutic response to immunomodulation with steroids is not impacted by sex-biased illness severity, which is consistent with the above observation of minimal sex differences in the inflammatory/immune response to acute sepsis.

**Figure 5.**
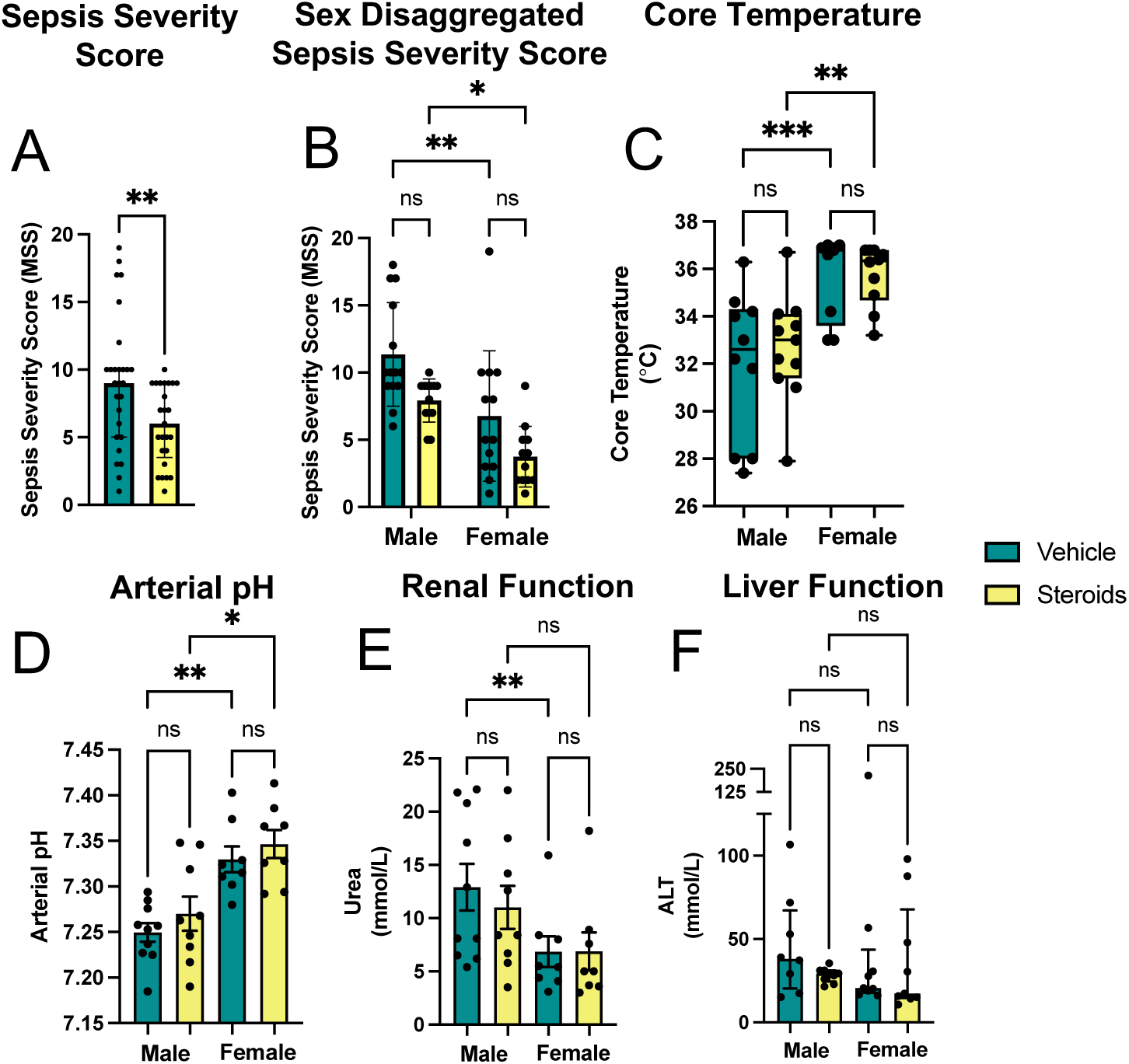
-Therapeutic targeting of immune-mediated tolerance with hydrocortisone does not impact sex-biased sepsis severity. (A) Murine sepsis severity index was quantified 6 hours post infection in both male and female mice that were treated with either steroid therapy (N=27, hydrocortisone succinate, 4 mg/kg, ip) or vehicle (N=25, PBS) followed by polymicrobial sepsis from intraperitoneal injection of donor cecal contents (1 mg/g). Sex disaggregated analyses were performed of steroid-treated males (N=13) and females (N=12), and vehicle treated males (N=14) and females (N=13) of (B) illness severity (MSSI), (C) core (rectal) temperatures, (D) arterial blood pH, and biochemical measurements of (E) plasma urea, and (F) plasma alanine aminotransferase (ALT). Dots are individual mice, bars show medians +/-interquartile range. Data were analyzed using Mann-Whitney U test (A) or 2-way ANOVA (B-F), ***p<0.0001, **p<0.01, *p<0.05, ns=p>0.05.

### Sexual dimorphism of mitochondrial tolerance in acute sepsis

To further interrogate host response differences that may contribute to sex biased sepsis severity, we explored the transcriptomic response to sepsis between males and females. We performed bulk RNA sequencing on hepatocytes from male and female littermates at 6 hours post infection with fecal peritonitis, as well as naïve uninfected controls. Vast transcriptomic changes were noted in response to sepsis in both males and females, but interestingly, responses differed significantly between males versus females (Fig 6A-B). Principal component analysis revealed clear separation of transcriptome responses between septic and naïve animals with 77% of the variance explained by infection status (Fig 6B). However, the second principal component, explaining 15% of the transcriptome variance, was biological sex, with clear clustering of transcriptomic responses between males and females under both naïve and septic conditions (Fig 6B).

**Figure 6.**
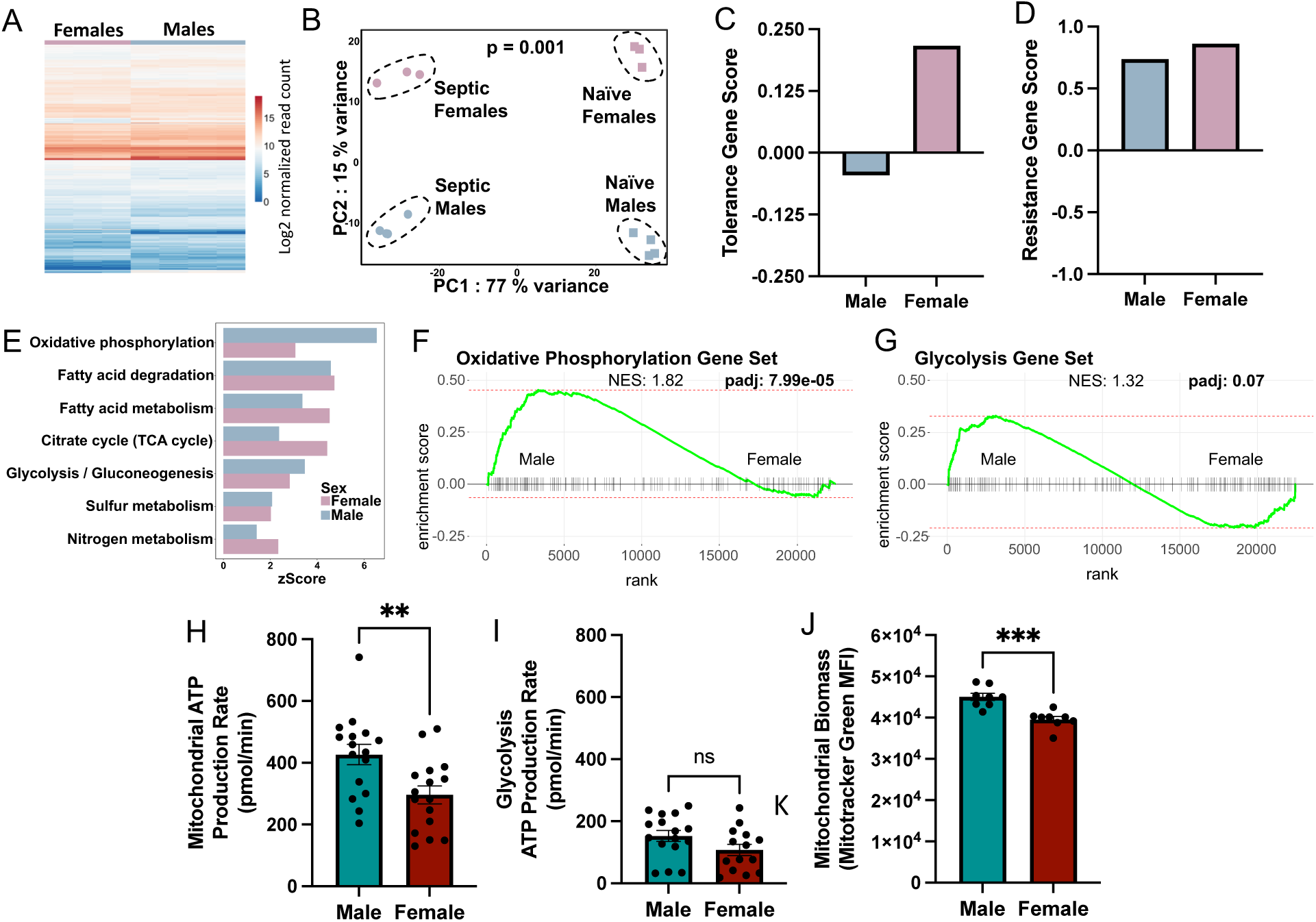
-Transcriptomic and cellular metabolic analyses of septic mice reveal sex-biased mitochondrial tolerance in sepsis. Bulk RNA sequencing of hepatocyte mRNA was performed in male (N=4) and female (N=3) mice at 6 hours post-infection with donor cecal contents (1 mg/g), as well as uninfected (native) control males (N=4) and females (N=3). (A) Heatmap showing log2-transformed read counts of differentially-expressed genes between septic males and females. (B) Principal components analysis of bulk RNA sequencing data from septic and uninfected (naïve) mice demonstrating clustering by sex and infection status. Analyzed by permutational analysis of variance (PERMANOVA), p=0.001. (C) Tolerance and (D) resistance gene scores were calculated from transcriptomic data of septic male and female mice. (E) KEGG pathway enrichment analysis (z score) of cellular metabolism pathways is shown between septic male and female mice. (E-F) Gene set enrichment analysis (GSEA) was performed between septic males and females showing (F) oxidative phosphorylation and (G) glycolysis gene sets. (H) Mitochondrial oxidative ATP production rate, and (I) glycolytic ATP production rate were measured by Seahorse XF analyzer, in hepatocytes isolated from septic male (N=16) and female (N=16) mice 6 hours post-infection with donor cecal contents (1 mg/g). (J) Total cellular mitochondrial biomass was measured by flow cytometry (MFI of Mitotracker Green stainin) in hepatocytes from septic male (N=8) and female (N=8) mice. For data in H-J, dots are individual mice, bars show mean +/- SEM, and data were analyzed using unpaired t-test, ***p<0.0001, **p<0.01, ns=p>0.05.

To identify gene expression pathways contributing to sex-biased sepsis severity, we began by evaluating gene regulation of pathways known to mediate infection resistance and tolerance between septic males and females. We utilized a recently developed tool described by Cohn et al.(*23*) that compares the magnitude of resistance versus tolerance gene programs in the mouse transcriptome by assigning a gene expression response score for host resistance gene program (R score) and host tolerance gene program (T score). We found that host resistance gene expression program (R score) was not different between septic males and females (Fig 6D), consistent with our observations above that pathogen burden was not different between sexes. In contrast, tolerance gene expression program (T score) trended higher in septic females compared to males, consistent with the higher illness severity (ie. impaired tolerance) observed in septic males (Fig 6C). Together with our data above, these findings further support the conclusion that differential illness severity between males and females is the result of sex-biased infection tolerance, rather than differential infection resistance.

Next, we explored sex differences in gene pathway enrichment between septic males and females using the KEGG database. KEGG pathway analysis revealed a number of expected sexually dimorphic pathways (e.g. steroid hormone biosynthesis), but interestingly 10 of the top 20 pathways that were differentially expressed between septic males and females were involved in cellular metabolism (Supplementary Fig 4). Amongst the KEGG pathways involved in cellular metabolism, the most prominent differences between males and females were observed in oxidative metabolism gene pathways (Fig 6E). To confirm these observations using an alternative approach to gene pathway analysis with gene set enrichment analysis (GSEA), we again observed prominent enrichment of oxidative phosphorylation gene set in septic males compared to septic females, whereas no sex-biased enrichment was observed for the glycolysis gene set (Fig 6F-G). To extend our findings from gene expression to cellular function, we performed a functional analysis of cell metabolism in hepatocytes isolated from septic males and females using Seahorse XF metabolic analyzer and Real-Time ATP assay. Consistent with our gene expression findings, oxidative (mitochondrial) ATP production rate was significantly higher in hepatocytes from septic males compared to septic females, whereas no differences were observed in glycolytic ATP production rates between sexes (Fig 6H-I). Of note, total cellular mitochondrial biomass measured by flow cytometry was slightly lower in female cells compared to male cells, but proportionally, this difference was minute compared to differences observed in gene expression and mitochondrial oxidative metabolism in males compared to females (Fig 6J). Taken together, these data demonstrate that male-biased sepsis severity is linked with impaired infection tolerance gene expression, and sexual dimorphism of cellular oxidative metabolism that collectively represents impaired mitochondrial tolerance response in septic males.

### Therapeutic correction of sex-biased illness severity in sepsis by targeting mitochondrial tolerance

Mitochondrial tolerance has emerged as a crucial determinant of illness severity and outcomes in bacterial and viral infections (*15*). Our observations of sex-biased infection tolerance linked to differential mitochondrial metabolism led us to examine the impact of mitochondrial tolerance in driving sexual dimorphism of sepsis severity. Multiple recent studies have established that tetracycline antibiotics, in particular doxycycline, are potentiators of mitochondrial tolerance in various mouse models of infection (*18, 19*). Therefore, we employed doxycycline treatment to test the hypothesis that sex-biased illness severity could be abrogated by therapeutically potentiating mitochondrial tolerance in males.

Following 3 daily treatments with doxycycline (1.75 mg/g i.p.) as previously described(*18*), male and female littermates were infected with a doxycyline-resistant strain of *E. coli* ST131. As previously reported, utilization of a doxycycline-resistant infection enables uncoupling of doxycycline antibiotic effect, ensuring that its impact on host mitochondrial tolerance and illness severity is independent of any impact on the growth and dissemination of the pathogen. We confirmed this with quantitative bacterial cultures of blood and tissues at 6 hours post-infection, confirming that *E. coli* ST131 infection was completely resistant to doxycycline treatment, with extensive pathogen dissemination in both vehicle and doxycycline treated groups (Fig 7A-C).

**Figure 7.**
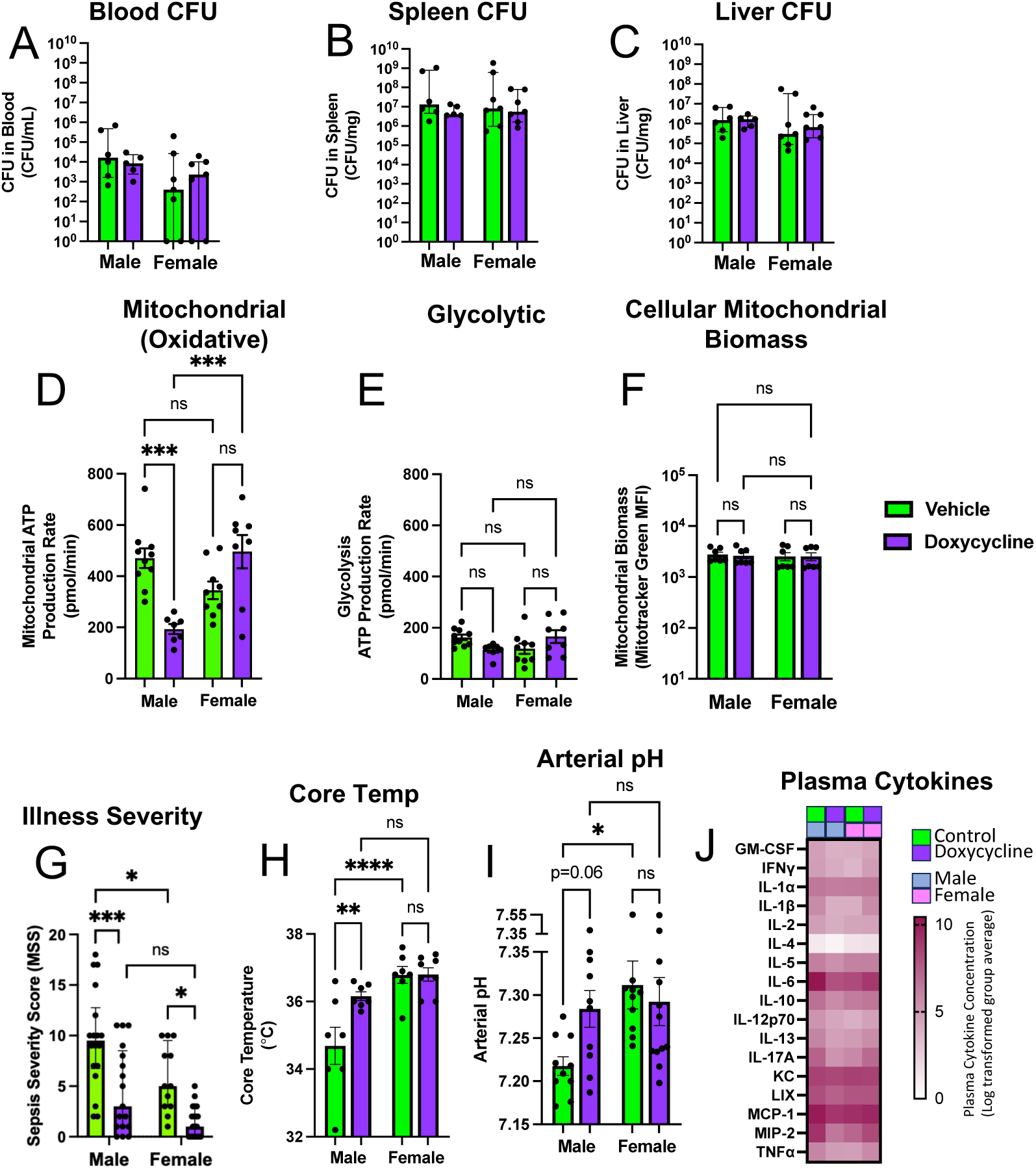
-Therapeutic potentiation of mitochondrial tolerance abrogates sex -biased sepsis severity. Male and female mice were treated with doxycycline (1.75 μg/g i.p. daily for 3 days) followed by infection with *E. coli* ST131 (2x10^7^ CFU, ip). Bacterial quantity (colony forming units, CFU) grown from (A) blood, (B) spleen, and (C) liver collected from vehicle-treated male (N=6) and female (N=7), and doxycycline-treated male (N=5) and female (N=7) harvested at 6 hours post-infection. (D) Mitochondrial oxidative ATP production rate, and (E) glycolytic ATP production rate were measured by Seahorse XF analyzer, and (J) total cellular mitochondrial biomass was measured by flow cytometry (MFI of Mitotracker Green staining) in hepatocytes isolated from vehicle-treated male (N=10) and female (N=9) and doxycycline-treated male (N=7) and female (N=8) mice 6 hours post-infection with donor cecal contents (1 mg/g). (G) Illness severity (MSSI), (H) core (rectal) temperature, (i) arterial pH, and (J) plasma cytokines were quantified in septic male and female mice treated with vehicle or doxycycline. Dots are individual mice, bars show median +/-interquartile range, and data were analyzed using 2-way ANOVA, ****p<0.00001, ***p<0.0001, **p<0.01, *p<0.05, ns=p>0.05.

Compared to vehicle-treated septic mice, treatment with doxycycline abrogated the sex difference in mitochondrial ATP production rates measured by Seahorse XF metabolic analyzer in hepatocytes from septic mice, with no impact on cellular glycolysis nor total mitochondrial biomass (Fig 7D-F). Modulation of mitochondrial tolerance with doxycycline treatment resulted in significantly reduced illness severity scores (MSSI) in septic males, with lesser improvement in females (Fig 7G). Male-biased illness severity observed in vehicle-treated septic mice was completely abrogated after doxycycline treatment (Fig 7G). In addition to behavioural illness severity, we also observed significant improvement of core temperature measurements as well as arterial pH in doxycycline-treated males, to levels that were indistinguishable from females (Fig 7H-I). Importantly, quantification of plasma cytokines/chemokine levels in vehicle and doxycycline-treated mice confirmed that doxycycline treatment did not alter the systemic inflammatory response in either males or females (Fig 7J). In summary, these data demonstrate that potentiation of mitochondrial tolerance with doxycycline results in reduced illness severity and organ dysfunction in septic males to the levels observed in females, yielding abrogation of sex-biased sepsis severity by targeting mitochondrial tolerance.

## DISCUSSION

In this study, we found that male-biased illness severity in bacterial sepsis is mediated by impaired mitochondrial tolerance, independent of infection resistance or canonical immune/inflammatory dysregulation. Therapeutic potentiation of mitochondrial tolerance with doxycycline neutralized sexual dimorphism of illness severity and organ dysfunction through a male-predominant treatment effect. These data reveal that biological sex is a fundamental determinant of illness severity and treatment-responsiveness in sepsis, and establish mitochondrial infection tolerance as a key mechanism driving sex-biased illness severity and treatment heterogeneity between males and females with sepsis.

Mitochondrial tolerance has emerged as an important determinant of sepsis outcomes in mouse models, mediated by shifts in cellular metabolism from oxidative phosphorylation towards glycolysis that mitigate the impact of oxygen and nutrient deprivation during infection (*19, 24*). Our interrogation of these mechanisms of mitochondrial tolerance revealed that males display impaired mitochondrial tolerance in sepsis, demonstrated by higher mitochondrial oxidative ATP production coupled with higher expression of genes involved in the oxidative phosphorylation pathway compared to females. The impact of sexually dimorphic mitochondrial tolerance towards sepsis severity was demonstrated by treatment doxycycline, which inhibits mitochondrial ribosomes and protein synthesis to potentiate tolerogenic metabolic shifts(*18*).

Doxycycline reduced mitochondrial ATP production rates, illness severity, and organ dysfunction in septic males but had minimal effect in septic females due to their superior baseline mitochondrial tolerance response in sepsis. Interestingly, estrogens have been reported as potent agonist of mitochondrial tolerance pathways in macrophages (*24*), consistent with our data from “four core genotype” transgenic mice that confirmed female-biased infection tolerance is driven by gonadal/hormonal sex phenotype, independent of sex-chromosomes. Together with mitochondrial tolerance, mammals have evolved multiple additional mechanisms of disease tolerance to counteract damage from infection and inflammation(*15*), including pathways to mitigate damage from reactive oxygen species(*25, 26*), tissue hypoxia(*27, 28*), glucose deprivation (*29, 30*), alterations in osmotic forces (*31*), as well as attenuation of inflammation in response to prolonged or repeated bacterial exposure (*32*). Whether additional cellular mechanisms of disease tolerance contribute to sex-biased sepsis outcomes will require further investigation in future studies.

The importance of biological sex towards immune function is well established (*33, 34*). Sex differences in immune-mediated diseases have been recognized for decades, including the observation that ∼80% of autoimmune diseases are female predominant (*33*). Thus, it was somewhat surprising that only modest differences in systemic inflammation and cellular immune responses were observed between septic male and female mice, and that sex-biased illness severity persisted after systemic anti-inflammatory (corticosteroid) treatment. While our investigation was focused on the systemic (circulating) immune response to peritonitis, prior studies have reported sex-based differences in organ- and pathogen-specific host defense. For example, in mouse models of bacterial peritonitis, higher mortality in male mice was linked with decreased density of surface TLR4 expression and dysregulated cytokine production by peritoneal macrophages in males, which was reversed by castration/androgen deprivation(*35, 36*). In mouse models of bacterial pneumonia, sex-biased immune responses varied depending on the infecting pathogen, with male-biased susceptibility seen in response to bacteria such as *A. baumanii* and *S. pneumoniae*, equivocal response to *S. aureus* and *K. pneumoniae*, or even female-biased immune defects and higher mortality in models of *P. aeruginosa* pneumonia (*37, 38*). Thus, the complex interplay between biological sex and immune mechanisms of host defense may have a nuanced impact on sepsis outcomes in a pathogen- and even organ-specific manner.

The demonstration that therapeutic targeting of mitochondrial tolerance yields a male-biased treatment response has important implications in preclinical, as well as clinical sepsis research. Sepsis research has been plagued by near-universal failure of promising preclinical therapeutic discoveries in mice to translate into efficacious treatment outcomes in human clinical trials (*13*). Our findings suggest that biological sex may represent a fundamental source of heterogeneity of treatment effect in sepsis that may contribute to this translational void in sepsis research. First, preclinical sepsis discoveries have historically relied on animal models using only (or largely) male subjects, potentially yielding inappropriate generalization of unrecognized sex-biased mechanisms. As a result, treatment may fail to show overall benefit when applied to both male and female participants. Indeed, a recent systematic review of animal sepsis studies reported an extreme paucity of reporting on sex-dependent treatment effects in preclinical sepsis research(*10*). Similarly, clinical trials of human sepsis often have male-biased enrollment(*39*), yet comprehensive analyses of clinical trial data between male and females are lacking. Recently, a post-hoc sex-disaggregated analysis of the ADRENAL trial, which found no benefit of hydrocortisone infusion on 90-day survival in patients with septic shock, reported sex-biased outcomes between males and females in which steroid therapy was associated with worse outcomes in females compared to males (more shock recurrence, longer duration of mechanical ventilation and ICU admission) (*40*). Thus, therapeutic strategies concluded to be non-beneficial when studied in mixed populations may in fact be efficacious, or even harmful, for one sex but not the other. Therefore, our findings support the inclusion of biological sex as a core variable to be evaluated in studies of pathogenesis mechanisms as well as treatment effects in pre-clinical as well as clinical trial investigations, both in future studies as well as potentially re-visiting prior with sex-disaggregated post-hoc analyses.

Our study has a number of limitations that warrant further research. First, our experiments focused on the acute phase of bacterial sepsis, as this is the most commonly studied phase of illness in preclinical sepsis models. However, it will be of great interest for future studies to investigate how biological sex impacts illness severity, organ dysfunction, and sepsis outcomes during the later subacute and even chronic phases of sepsis. Furthermore, it will be of interest to understand the impact of sex on long term sepsis outcomes in survivors including the post-sepsis syndrome of chronic organ dysfunction and long-term mortality risk that plagues many sepsis survivors. In addition, experiments in this study intentionally utilized a non-lethal model of sepsis, therefore it will be of interest for future studies to determine whether sex-based difference in mitochondrial tolerance, illness severity, organ dysfunction, and treatment effect of mitochondrial tolerance potentiation (doxycycline) persist in more severe infections, or whether there are threshold of severity beyond which sex-based differences are nullified. Notably, prior studies of mitochondrial tolerance potentiation with doxycycline in male mice did show a survival benefit in lethal models of infection(*18*), but whether this survival benefit extends to females is unknown. Finally, it will be of interest for further work to explore other organ- and pathogen-causes of sepsis such as pneumonia models, as this may uncover additional nuances of sex-based illness severity and organ dysfunction.

## MATERIALS AND METHODS

### Animal husbandry and treatments

Animal protocols performed were all approved by the University of Calgary Animal Care Committee and followed in accordance with the Canadian Guidelines for Animal Research (Protocol #: AC19-0139). All experimental mice used were of a C57BL/6 genetic background and aged 8-12 weeks, bred and maintained under standard specific pathogen free conditions at the University of Calgary Animal Resource Centre. Transgenic Four Core Genotype mouse embryos (B6.Cg-Tg(Sry)2Ei *Sry^dl1Rlb^* T(XTmsb4x-Hccs;Y)1Dto/ArnoJ) were purchased from The Jackson Laboratory (strain #010905) and implanted in C57BL/6 females, and hemizygous transgenic males from the F1 generation (XYSry^dl1Rlb^Tg(Sry)2Ei) were bred with wild-type C57BL/6 females yielding the 4 core genotypes (XX wildtype (XX with ovaries), XY *Sry^dl1Rlb^* (XY with ovaries), XX Tg(Sry)2Ei (XX with testes), and XY Sry^dl1Rlb^ (XY with testes).

Genotype confirmation of every 4CG mouse used in this study was performed by PCR from ear samples as per Jax protocol 5990 V1.0. In some experiments, mice were treated with hydrocortisone succinate (Pfizer, 4 mg/kg, i.p.) prior to infection, doxycycline (Millipore-Sigma, 1.75 mg/g, i.p.) as previously described(*18*) daily for 3 days prior to infection, or vehicle controls (equivalent volumes of sterile PBS, i.p.).

### Fecal-induced peritonitis (FIP) model of sepsis

The fecal slurry preparation was adapted from previously published protocol (NPSP paper). Donor C57BL/6 mice at 8-12 weeks of age were euthanized by cervical dislocation under inhaled isofluorane anesthetic, and cecal contents were collected and homogenized in phosphate buffered saline with a 5 mL syringe to break up the contents. The slurry was filtered using a 100 μM cell strainer to remove large particles and centrifuged at 3,000 g for 25 min at 4 °C. The pellet was then resuspended in phosphate buffered saline a final concentration of 100 mg/mL. Mice were given a single intraperitoneal (IP) injection of 1 mg/g fecal slurry (in a total volume of 200 μL) into the right lower abdominal quadrant. Control mice received an IP injection of an equivalent volume of PBS vehicle.

### Monomicrobial model of sepsis using *E. coli* ST131

*Escherichia coli* ST131 was grown from 2μL of frozen glycerol stock that was thawed and combined with 10mL of Lysogeny Broth (LB) in a shaking incubator at 37°C overnight. 5mL of LB was combined with 1mL of overnight culture and placed back in the shaking incubator for 2 hours at 37°C to achieve log phase growth. 1 mL of this log-phase subculture was centrifuged at 8000rcf for 5 minutes and the supernatant was removed. The remaining contents were vortexed with 1 mL of PBS and diluted to 5x10^7^ CFU per injection volume (200μL). Mice were infected by intraperitoneal injection with 5x10^7^ CFU (in 200 μL PBS), whereas controls were injected with 200 μL of PBS vehicle.

### Bacterial quantification in tissues

Mice were anesthetized with 5% isoflurane, blood was collected by cardiac puncture, and animals were euthanized by cervical dislocation. Organs were harvested, weighed, and homogenized in 1 mL of sterile PBS using a TissueLyser II (Qiagen). Tissue homogenates and whole blood were diluted in BHI broth, and 10-fold dilutions were spotted on BHI agar (for fecal peritonitis experiments) or LB agar (for *E. coli* infection experiments) in triplicate. Agar plates were incubated overnight at 37°C under aerobic (and anaerobic in some experiments) conditions and bacterial colonies were counted and CFU/g of tissue or per mL of blood were calculated.

### Illness severity measurements

Core temperature was measured in infected mice using a rectal thermometer. Physiologic illness severity was quantified using the murine sepsis severity index (MSSI), a well-established and validated scoring system for quantitative analysis of sepsis severity in mouse models (*20*). Mice were independently observed at timepoints after infection (indicated in figure legends) and scored as described by Shrum *et al.*(*20*) on a scale of 0-4 for seven clinical variables including appearance, level of consciousness, activity, response to stimulus, eyes, respiration rate, and respiration quality. The total score is reported.

### Mass Cytometry Analysis

Immune cell profiling was performed using mass cytometry on cryo-preserved whole blood. Mice we anesthetized with 5% isofluorane and whole blood was collected via cardiac puncture into heparin containing tubes. 300μL of whole blood mixed with 500 μL PROT1 Proteomic Stabilizer (SmartTube, Inc) and stored at -80°C for batched analysis. For preparation of single cell suspensions, cryopreserved whole blood as thawed to room temperature and red blood cell lysis was performed using Thaw-Lyse Buffer (SmartTube, Inc.) following manufacturer’s instructions. The single cell suspension was then passed through a 100 μM cell strainer to remove large aggregates of cells or debris. Cells were then fixed, permeabilized, and barcoded using the Cell-ID 20-Plex Pd Barcoding Kit (Standard Biotools) following manufacturer instructions. Fixed and barcoded cells were then pooled and stained with a panel of antibodies (Supplementary Table 1) followed by DNA staining with Cell-ID Intercalator-Ir (Standard Biotools) overnight. Pooled samples were then acquired on a Helios CyTOFII mass cytometer (Standard Biotools). Acquired samples were debarcoded and imported into Cytobank (Beckman Coulter) for manual gating of individual cell populations (Supplementary Figure 3). For unsupervised analysis, CD45+ cells manually gated in Cytobank were exported to R for further analysis as previously described(*41*).

### Bulk RNA-Sequencing and Analysis

Mouse liver tissue was harvested and then stored in RNAlater at 4°C overnight and transferred to -80°C freezer. Thawed RNAlater-preserved liver tissue was homogenized by bead-beating, followed by RNA separation with trizol and chloroform, and RNA isolation using the RNeasy Mini kit (Qiagen) according to the manufacturer’s instructions. Total RNA (200 μg) was used for library preparation with the NEBNext Ultra II Directional RNA library prep kit (New England Biolabs), and mRNA was enriched with the NEBNext Poly(A) mRNA Magnetic Isolation Module (New England Biolabs) according to the manufacturer’s instructions. 100bp paired-end sequencing was conducted using the NextSeq 2000 platform with a target read depth of 20-25M read-pairs per sample. As controls, we included External RNA Controls Consortium spike-ins (*42*) in 2 samples confirm fidelity of sequencing. RNA-Seq analyses were carried out in R (v4.5.0). Raw sequencing reads were pseudo-aligned to the *Mus musculus* reference transcriptome GRCm39 using *kallisto* (v0.46.1). Low-abundance transcripts were filtered out if they were present in less than two samples. Differential gene expression analysis was performed using *DESeq2* (v1.48.1). Gene functions were annotated using *biomaRt* (v2.64.0) to access the Ensembl database (*43*). Tolerance and resistance gene expression scores were calculated as described by Cohn *et al.*(*23*). KEGG pathway enrichment analysis was conducted using *cluterProfiler* (v4.16.0) and gene set enrichment analysis was conducted using *fgsea* (v1.34.0).

### Biochemical measurements and Luminex Immunoassay

Mouse blood was collected from anesthetized animals by cardiac puncture into heparin-coated syringes. One hundred μL of fresh whole blood was loaded into an EPOC blood analysis system (Siemens Healthcare) cartridge for measurement of blood pH, blood glucose, and urea. The remaining blood was centrifuged at 2,000g for 10min at 4 °C, after which the plasma was separated and stored at −80°C for further analyses. Cytokine measurements were performed on thawed plasma samples using Luminex Immunoassay Mouse High Sensitivity T-Cell 18-Plex Discovery Assay (MDHSTC18) (performed by Eve Technologies, Calgary, Canada). Alanine aminotransferase (ALT) levels in thawed plasma was determined by using Mouse ALT ELISA Kit (ab282882, Abcam) as per manufacture’s protocol.

### Cell metabolism and mitochondrial analyses

Livers were harvested from mice, minced with scissors, and incubated at 37°C for 30 mins in a shaking incubator in an enzymatic tissue dissociation buffer (RPMI containing 10% heat-inactivated horse serum, 100 mM HEPES, collagenase VIII 0.5 mg/mL, DNase I 5 U/mL).

Digested liver samples were then passed sequentially through 100 μm then 70 μm cell strainers, washed, and underwent differential centrifugation at 60g for 5 mins at 4°C (no brake) to pellet large liver parenchymal cells (primarily hepatocytes). The pellet was collected and residual RBCs were lysed by incubation in ACK lysis buffer for 10 mins. The isolated cells were counted using a hemocytometer, and seeded into Seahorse XF cell culture microplates at 1.5 × 10^5^ cells per well. Sensor cartridges were hydrated overnight in Seahorse XF Calibrant at 37 °C without CO₂ and calibrated in the Seahorse XF Pro Analyzer (Agilent Technologies). Cells were washed once with prewarmed XF DMEM assay medium supplemented with 10 mM glucose, 2 mM L-glutamine, and 1 mM sodium pyruvate, and then equilibrated at 37 °C without CO₂ for 45-60 minutes. The real-time ATP Rate Assay protocol was performed as per the manufacturer’s instructions (Agilent Technologies), comprising three baseline measurement cycles followed by sequential injections of oligomycin (1 µM) and rotenone/antimycin A (0.5 µM each), followed by three measurement cycles. Oxygen consumption and extracellular acidification rates were recorded using Wave software (Agilent Technologies), and mitochondrial versus glycolytic ATP production rates were calculated with the XF Real-Time ATP Rate Report Generator (Agilent Technologies). Data were normalized to cell number and reported as ATP production rate in pmol/min. In addition, total mitochondrial biomass was measured in isolated hepatocytes by flow cytometry. Briefly, isolated cells were incubated with Zombie Aqua viability dye (Thermo Fisher Scientific) followed by Fc receptor blocking using anti-mouse CD16/CD32 (clone 93; BioLegend). Cells were then surface stained for 20 min at 4°C with the following fluorochrome-conjugated antibodies: Anti-CD45 PerCP (clone 30-F11; BioLegend), Anti-CD11b APC/Cy7 (clone M1/70; BD Biosciences). After surface staining, cells were incubated with MitoTracker Green FM (100 nM; Invitrogen) for 15 min at 37°C. Data acquisition was carried out on a Cytek Aurora spectral cytometer, and data analysis was performed using FlowJo v10.

Non-leukocyte parenchymal cells (primarily hepatocytes) were gated as live, CD45^-^, CD11b^-^ events, and mitochondrial biomass was quantified by the median fluorescence intensity (MFI) of MitoTracker Green within this gate. Fluorescence Minus One (FMO) controls were used to set gates for surface markers.

### Statistical Analyses

For all experiments, biological replicates consisted of individual mice and the number of replicates (n) is indicated in figure legends. Normally distributed data are show as mean ± standard error of the mean (SEM) and compared using unpaired T-test (to compare 2 groups) or one-way ANOVA followed by Tukey’s post-hoc (for >2 groups). Non-normally distributed data are shown as median with interquartile range and compared using Mann-Whitney U test (for 2 groups), or Kruskal-Wallis followed by Dunn’s multiple comparisons test (for >2 groups). Data with two independent variables (*e.g.*, males versus females and treatment versus vehicle) were analyzed by 2-way ANOVA with Tukey’s multiple comparisons test. For statistical comparisons of transcriptomic clustering between experimental groups, data were analyzed using permutational ANOVA (PERMANOVA) on the calculated Bray-Curtis distances. KEGG pathway analysis within RNA-sequencing datasets utilized the hypergeometric test with Bejamini-Hochberg correction for multiple comparisons, using the R package *vegan* (v2.6-10). Gene set enrichment analysis (GSEA) was performed using the R package *fgsea* (v1.34.0) which uses an adaptive multi-level split Monte-Carlo scheme to estimate p-values. Data analysis and visualization were performed using R version 4.5.0 for RNA sequencing analysis, and GraphPad Prism® version 10.4 (GraphPad Software, Inc).

## LIST OF SUPPLEMENTARY MATERIALS

Supplementary Table 1

Supplementary Figure 1-4

## ACKNOWLEDGEMENTS

The authors would like to thank the Nicole Perkins Microbial Core at the Cumming School of Medicine for assistance with mass cytometry, as well as the Cumming School of Medicine Genomic Core for assistance with RNA sequencing experiments.

## AUTHOR CONTRIBUTIONS

Conceptualization - BD, and BM Methodology - BD, KS, AW, JS, DC, NAC, IY, and BM Investigation - BD, KS, AW, JS, DC, NAC, IY, and BM Analyses - BD, KS, AW, JS, DC, NAC, IY, and BM Supervision - BM Writing – original draft - BD and BM Writing – review and editing - BD, KS, AW, JS, DC, NAC, IY, and BM

## DATA AND MATERIALS AVAILABILITY

RNA sequencing data is available in the National Institutes of Health Gene Expression Omnibus under the accession number GSE303231. Analysis pipelines utilized publicly available code as noted in the materials and methods section, with additional details available upon request. All other data is available in the main text or supplementary materials.

