## Supplementary Materials for "Differential disease tolerance mediates sex-biased illness severity in sepsis"

For

List of Supplementary Materials

Supplementary Table 1

Supplementary Figures 1 – 4

Supplementary Table 1 – Mass Cytometry Antibody Panel Details

| **Target** | **Clone** | **Metal Label** | **Catalogue Number** | **Supplier** |
|---|---|---|---|---|
| CD64 | X54-5/7.1 | Cd111Di | 139301 | Biolegend* |
| CD25 | 2A3 | Cd112Di | 3169003B | Biolegend* |
| CD43 | eBioR2/60 | Cd113Di | 14-0431-82 | Thermo Fisher* |
| IgM | RMM-1 | Cd114Di | 406527 | Biolegend* |
| IgD | 11-26c.2a | Cd116Di | 405737 | Biolegend* |
| Ly-6G | 1A8 | Pr141Di | 3141008B | Standard Biotools |
| IL-17RA | 657603 | Nd142Di | MAB4481 | R&D* |
| CD11b (Mac-1) | ICRF44 | Nd143Di | 3167011B | Standard Biotools |
| CD45R/B220 | RA3-6B2 | Nd144Di | 3144011B | Standard Biotools |
| CD4 | RPA-T4 | Nd145Di | 3145001B | Standard Biotools |
| F4/80 | BM8 | Nd146Di | 3146008B | Standard Biotools |
| CXCR2 | SA044G4 | Sm147Di | 149302 | Biolegend* |
| CD279 (PD-1) | EH12.2H7 | Nd148Di | 3174020B | Standard Biotools |
| CD23 | B3B4 | Sm149Di | 101625 | Biolegend* |
| Ly-6C | HK1.4 | Nd150Di | 3150010B | Standard Biotools |
| CD49d (Integrin α4) | R1-2 | Eu151Di | 3151016B | Standard Biotools |
| CD3 epsilon | 145-2C11 | Sm152Di | 3152004B | Standard Biotools |
| CD274 (PD-L1) | 10F.9G2 | Eu153Di | 3153016B | Standard Biotools |
| CD11a | M17/4 | Sm154Di | 101101 | Biolegend* |
| RORγt | B2D | Gd155Di | 3159019B | Standard Biotools |
| CCR2 | 475301R | Gd156Di | MAB55381R-100 | R&D |
| CD80/B7-1 | 16-10A1 | Gd158Di | 92J023158 | Standard Biotools |
| CD184 (CXCR4) | 12G5 | Tb159Di | 3156029B | Standard Biotools |
| CD62L (L-selectin) | MEL-14 | Gd160Di | 3160008B | Standard Biotools |
| CD40 | HM40-3 | Dy161Di | 3161020B | Standard Biotools |
| CD1d | 1B1 | Dy162Di | 3162020B | Standard Biotools |
| CD170 (Siglec-F) | S17007L | Dy163Di | 92J017163 | Standard Biotools |
| CX3CR1 | 2A9-1 | Dy164Di | 3172017B | Standard Biotools |
| FoxP3 | 259D/C7 | Ho165Di | 3159028A | Standard Biotools |
| CD19 | HIB19 | Er166Di | 3142001B | Standard Biotools |
| Gata3 | TWAJ | Er167Di | 3167007A | Standard Biotools |
| CD8a | RPA-T8 | Er168Di | 3146001B | Standard Biotools |
| Sca-1 | D7 | Tm169Di | 3169015B | Standard Biotools |
| CD161 (NK1.1) | PK136 | Er170Di | 3170002B | Standard Biotools |
| CD44 | IM7 | Yb171Di | 3171003B | Standard Biotools |
| CD86 | GL1 | Yb172Di | 3172016B | Standard Biotools |

| | | | | |
|---|---|---|---|---|
| CD69 | H!.2F3 | Yb173Di | 3113002 | Standard Biotools |
| MHC Class II | M5/114.15.2 | Yb174Di | 3174003B | Standard Biotools |
| CD127/IL-7R alpha | A7R34 | Lu175Di | 3175006B | Standard Biotools |
| CCR3 | J073E5 | Yb176Di | Novus Biologicals | MAB1551-100 |
| CD11c | Bu15 | Bi209Di | 3147008B | Standard Biotools |
| CD45 | HI30 | Y89Di | 3089003B | Standard Biotools |

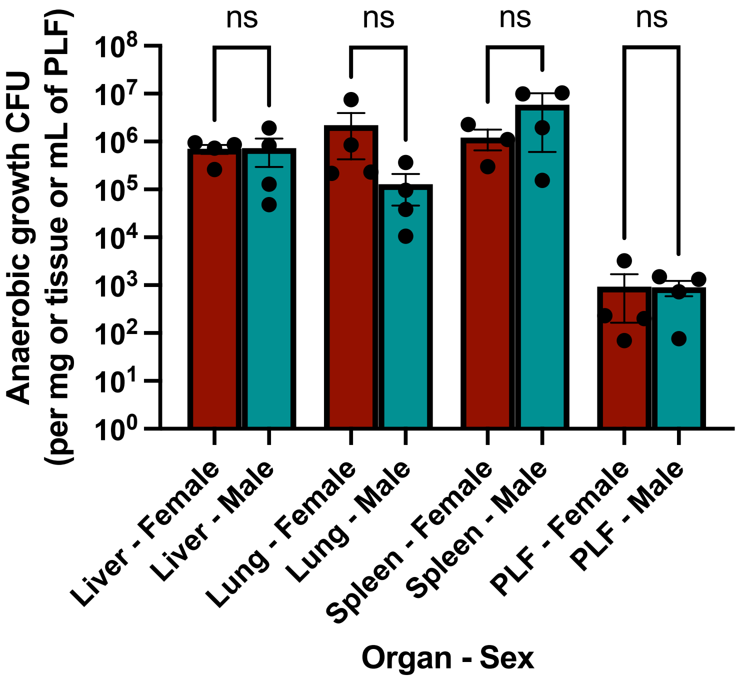


**Supplementary Figure 1 – No difference in anaerobic bacterial growth in organs between male and female mice after polymicrobial sepsis.** Male (N=4) and female (N=4) age-matched 8-12 week old littermates were infected intraperitoneally with donor cecal contents (1 mg/kg) to induce polymicrobial sepsis. At 6 hours post-infection, bacterial CFU were quantified in peritoneal fluid (PLF), spleen, liver, and lung by growth on BHI agar plates under anaerobic conditions. Dots are individual mice, bars show medians +/- interquartile range. Data were analyzed using Mann-Whitney U test, ns=p>0.05.

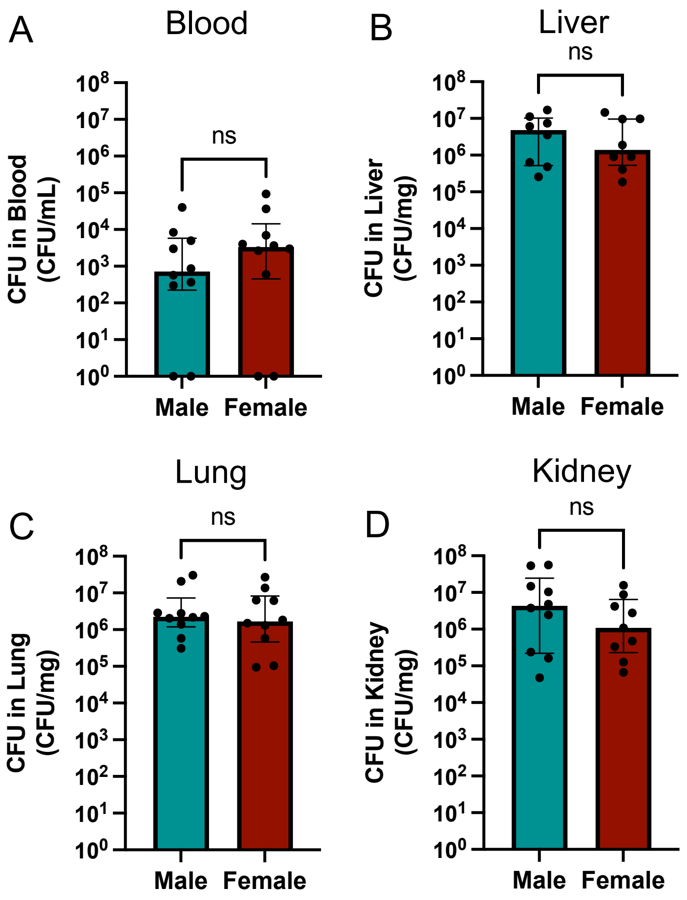


**Supplementary Figure 2 - No difference in bacterial burden in between male and female mice in mono-bacterial sepsis.** Male (N=10) and female (N=10) age-matched 8-12 week old littermates were infected intraperitoneally with *E. coli* ($2x10^7$ CFU, ip) to induce mono-bacterial sepsis. At 6 hours post-infection, bacterial CFU were quantified in (A) blood, (B) liver, (C) lung, and (D) kidney. Dots are individual mice, bars show medians +/- interquartile range. Data were analyzed using Mann-Whitney U test, ns=p>0.05.

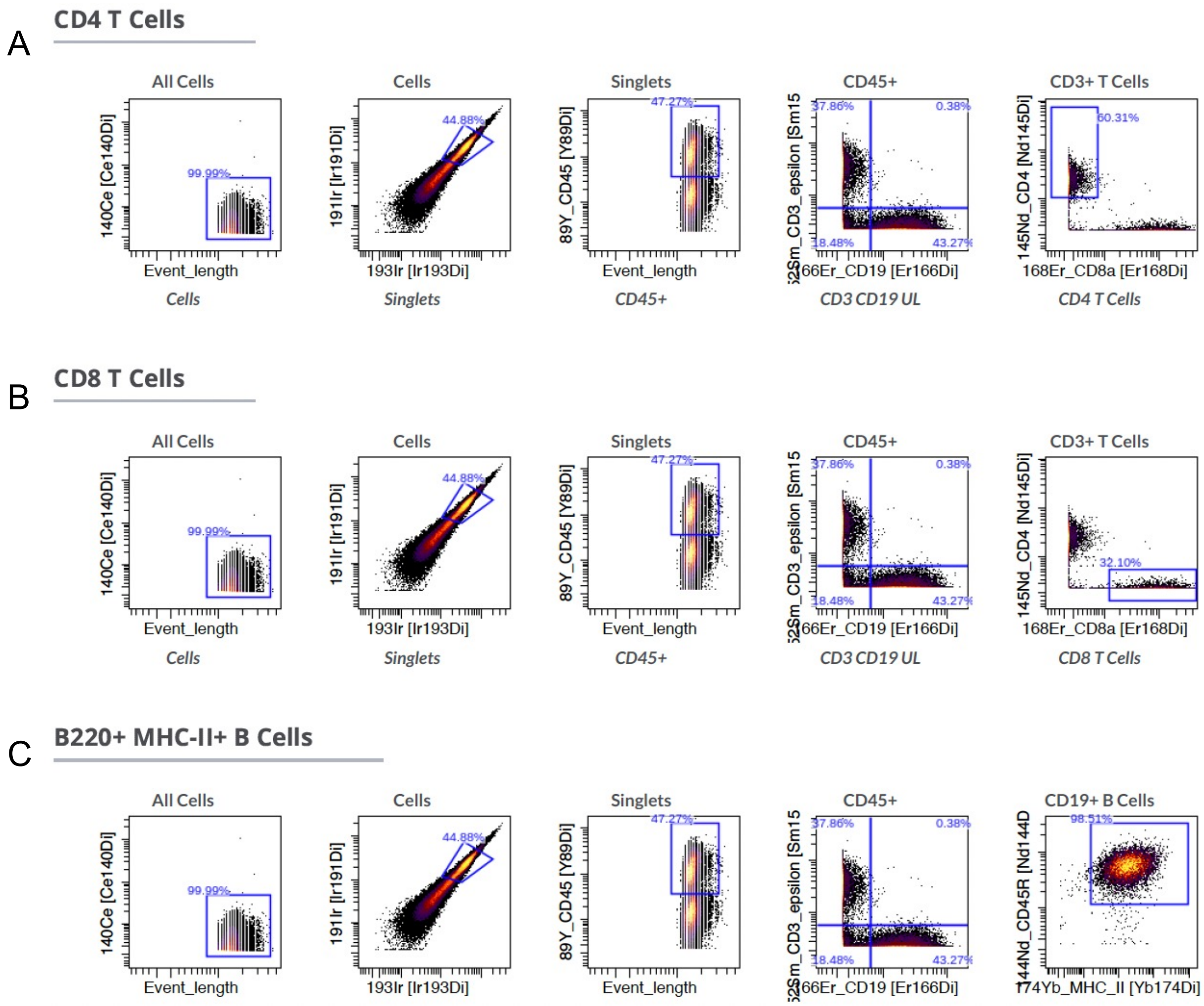


Continued on next page

D

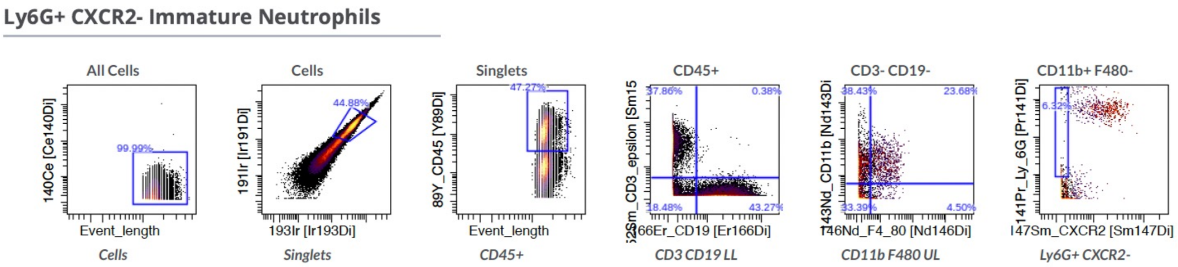


E

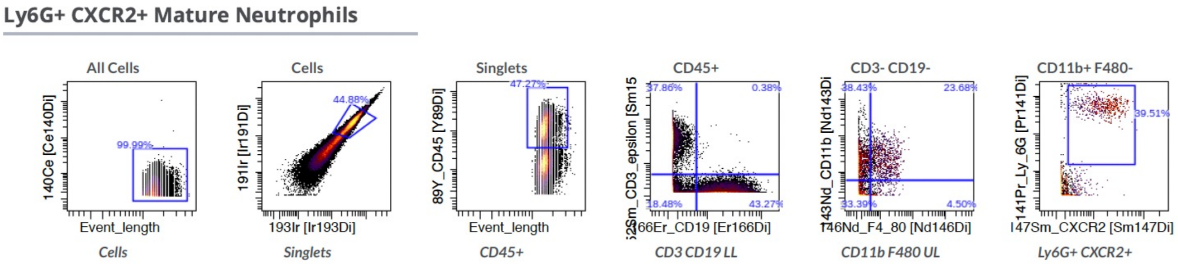


F

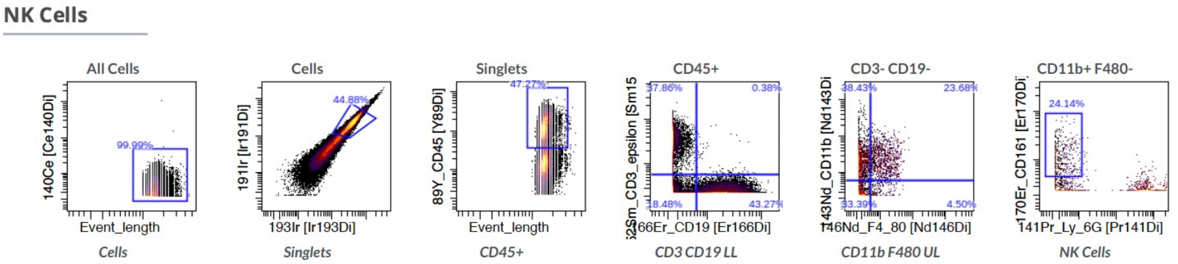


Continued on next page

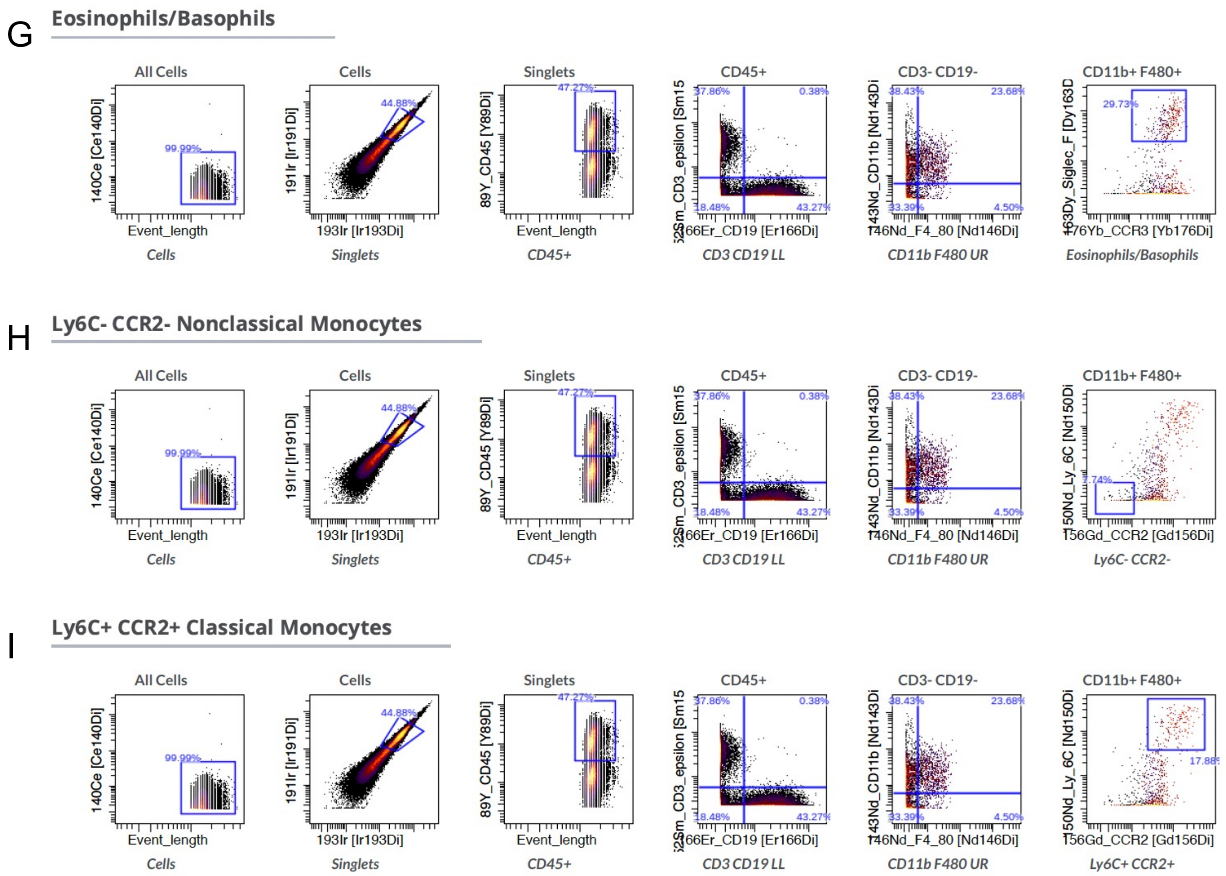


**Supplementary Figure 3 – Mass cytometry gating strategy.** Manual gating strategies for (A) CD4 T cells, (B) CD8 T cells, (C) B cells, (D) Immature neutrophils, (E) mature neutrophils, (F) NK cells, (G) Eosinophils, (H) non-classical monocytes, (I) classical monocytes.

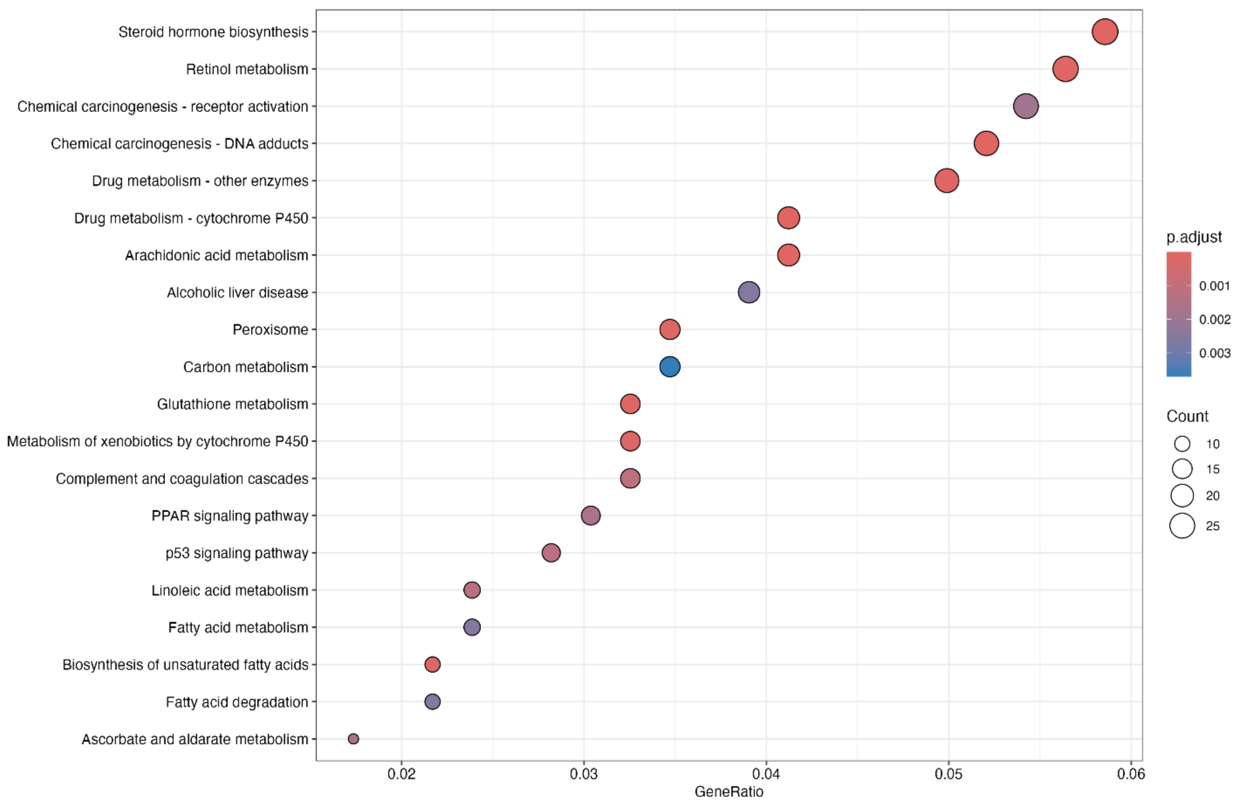


**Supplementary Figure 4 - KEGG pathway enrichment analysis showing top 20 differentially-enriched pathways between septic males and females.** X-axis represents the proportion of genes in the pathway that are differentially expressed between septic males and females. The size of the dots represents the number of genes in the pathway that are differentially expressed. Results are coloured by the adjusted p-value.
